# Topological data analysis reveals a core heteroblastic program embedded in leaves of grapevine and maracuyá

**DOI:** 10.1101/2023.07.14.549010

**Authors:** Sarah Percival, Daniel H. Chitwood, Aman Y. Husbands

**Affiliations:** Department of Computational Mathematics, Science & Engineering, Michigan State University, East Lansing MI 48823; Department of Horticulture, Michigan State University, Michigan State University, East Lansing MI 48823; Department of Biology, University of Pennsylvania, Philadelphia PA 19104 USA; Epigenetics Institute, University of Pennsylvania, Philadelphia PA 19104 USA

**Keywords:** topological data analysis, Mapper algorithm, heteroblasty, leaf development, evolution

## Abstract

Leaves have a single shape. However, embedded in that shape are a multitude of latent shapes arising from evolutionary, developmental, environmental, and other effects. These confounded effects manifest at distinct developmental time points and evolve at different tempos. Here, revisiting datasets comprised of thousands of leaves of vining grapevine (Vitaceae) and maracuyá (Passifloraceae) species, we apply a technique from the mathematical field of topological data analysis to comparatively visualize the structure of heteroblastic effects on leaf shape in each group. Consistent with a morphologically closer relationship, members of the grapevine dataset possess a strong core heteroblasty program with little deviation between species. Remarkably, we found that most members of the maracuyá family also share a core heteroblasty program despite dramatic species-to-species leaf shape differences. This conservation was not detected using traditional analyses. We also identify two morphotypes of maracuyá that deviate from the core structure, suggesting the evolution of new heteroblastic properties in this phylogenetically related sub-group. Our findings illustrate how topological data analysis can be used to disentangle previously confounded developmental and evolutionary effects to visualize latent shapes and hidden relationships, even ones embedded in complex, high-dimensional datasets.

## INTRODUCTION

Leaf shape is dynamic. Rather than viewing the ways it changes in response to evolutionary, developmental, and environmental forces as facets of a single form, we partition these effects separately from each other. This was not always the case, and in early philosophical conceptualizations of the plant form, evolutionary, developmental, and environmental responses flowed seamlessly with each other, focusing on the organismal form. Before Darwin, the idea of gradual change as the foundation of evolutionary thinking was elaborated by Goethe, focusing on the metameric, serial homology between leaves and other plant organs <u>(Friedman and Diggle 2011)</u>. In describing changes to mature leaf shape across sequential nodes (what we now call heteroblasty) as well as more dramatic metamorphoses of lateral organs, Goethe <u>(Goethe 2000)</u> declared that the ideal leaf is mutable and its only constant is change itself: “*daß in demjenigen Organ der Pflanze, welches wir als Blatt gewöhnlich anzusprechen pflegen, der wahre Proteus verborgen liege*” (“that in the organ of the plant which we are accustomed to calling the leaf, the true Proteus lies hidden”). Experiments by Hales pricking a grid of points in a young fig leaf and measuring vertical and horizontal displacement as it expanded determined that, not only does leaf shape change across successive nodes, but that the shape of each individual leaf constantly changes during its development: “By observing the difference of the progressive and lateral motions of these points in different leaves, that were of very different lengths in proportion to their breadths.” <u>(Hales 1969)</u>. The environment, too, modulates leaf shape, through heteroblastic and ontogenetic processes. Evolution and the environment thus act on multiple developmental processes, including ontogeny and heteroblasty, to generate inter-species, intra-species, intra-individual, and intra-leaf variation in shape <u>(Diggle 2002)</u>. Discerning the relative contributions of these developmental, environmental, and evolutionary forces in leaf morphogenesis remains a key challenge.

We think of leaf shape geometrically, in terms of the relative size and distance of features to each other <u>(Viscosi and Cardini 2011)</u>. Lobes, serrations, and leaf dimensions are all examples of this geometry. Landmarks – homologous points found on every leaf – allow geometric features to be quantified <u>(Bookstein 1992)</u>. For instance, Generalized Procrustes Analysis (GPA) can be used to superimpose leaves on each other <u>(Gower 1975)</u> using transposition, scaling, and rotation to minimize the distance of landmarks on every leaf to each other. Once superimposed, *x*- and *y*- coordinate values can be modeled and analyzed statistically. Using these methods on grapevine and maracuyá leaves, evolutionary (<u>Chitwood, Klein, et al. 2016; Chitwood and Otoni 2017b)</u>, developmental <u>(Bryson et al. 2020; Chitwood and Otoni 2017a)</u>, and environmental effects <u>(Baumgartner et al. 2020; Chitwood et al. 2021; Chitwood, Rundell, et al. 2016)</u> can be studied. Ultimately these approaches are statistical and treat each leaf as a separate sample. Population parameters like mean and standard deviation of coordinate values are calculated as summaries and dimension reduction is used to efficiently analyze multivariate data. However, each leaf remains a separate entity from every other, and only the statistical parameters at a population level are modeled.

Just as features within a single leaf have a relative distance to each other, each leaf has a relative distance to other leaves, based on their overall similarity. By computing the correlation distance between each leaf shape, a matrix of all the distances from each leaf to every other leaf can be generated. This distance matrix itself has a visualizable structure: in effect, a “shape of shapes”. Importantly, this structure changes depending on the mathematically defined perspective from which we view it. Topological data analysis is a mathematical field that measures the structure of data by its topology – connected components, loops, voids and other robust features that only change by tearing or detachment (see <u>(Munch 2017)</u> for a brief overview and <u>(Dey and Wang 2022)</u> for a more thorough treatment). The Mapper algorithm <u>(Singh et al. 2007)</u> is a method from topological data analysis that visualizes data structures as graphs or networks. The graph structure is primarily determined by a lens function: a real number value assigned to each data point that determines the data structure from a mathematically defined perspective. Mapper has been used to visualize biological data structures from functional- and hypothesis-driven perspectives, including the discovery of cancer-associated genes <u>(Rabadán et al. 2020)</u> and discerning developmental transitions in single-cell transcriptomic studies <u>(Rizvi et al. 2017)</u>.

Here, we use topological data analysis to identify developmental and evolutionary relationships hidden within datasets of leaf shape in grapevine (Vitaceae) and maracuyá (Passifloraceae). Both families are noted for the disparate leaf shapes that characterize different species, with maracuyá in particular displaying extreme differences in leaf shape thought to arise from selective pressures of *Heliconius* butterflies laying eggs on its leaves (**Fig. 1**; <u>Dell’Agli et al. 2016</u>). Using the relative node position of leaves within the shoot as a lens function, we comparatively visualized the developmental progression of leaf shape in each family as a Mapper graph. Our analyses suggest that leaves of grapevine species progress through a nearly identical heteroblastic program. Surprisingly, heteroblastic progression in maracuyá species is also strongly conserved despite the strikingly different leaf shapes in this charismatic family. This suggests the acquisition of new morphologies in groups such as maracuyá may result from modifications resting on top of deeply conserved developmental programs, like heteroblasty, rather than by alterations to the heteroblasty program itself. Our analyses also identify two interesting exceptions to this conservation which cluster in the *Decaloba* subgroup of maracuyá. These species appear to have modified their heteroblasty program, but only at nodes near the middle of the shoot, creating a “reverse hourglass” effect. Taken together, our findings illustrate the power of topological data analysis to isolate conserved developmental relationships hidden within large, high-dimensional datasets, with implications for the development of predictive models of complex phenotypes.

**Figure 1.**
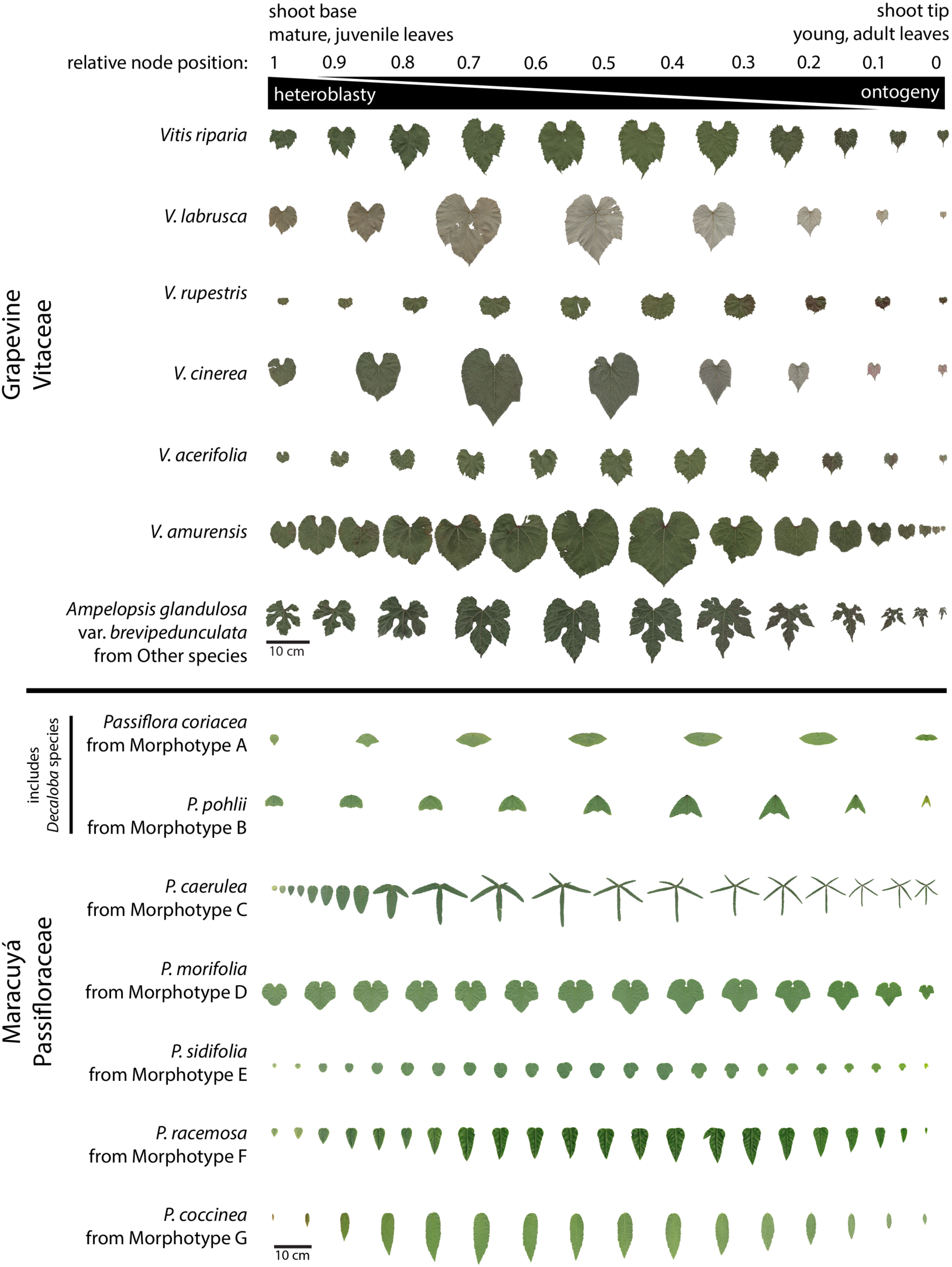
Leaves of grapevine and maracuyá separated by relative node position. Leaves are arranged from shoot base (mature, juvenile) to shoot tip (young, adult). Relative node positions are then assigned for each leaf with 1 being base and 0 being tip. Heteroblasty is evident for all species but is particularly striking in the maracuyá group. Species-to-species differences in leaf morphology is also more dramatic in the maracuyá group. Scale bar = 10cm.

## MATERIALS AND METHODS

### Plant materials

Grapevine data is previously described in Chitwood et al. <u>(Chitwood et al. 2021; Chitwood, Klein, et al. 2016; Chitwood, Rundell, et al. 2016</u>). Original leaf scans <u>(Chitwood et al. 2020)</u> and landmark data <u>(Chitwood 2020)</u> are both publicly available. More than 8,400 leaves were collected from up to 11 species and four hybrids representing 208 vines over four different years keeping track of the node, counting from the growing tip of the vine, they were collected from. The species and hybrids are as follows: *Vitis riparia*, *V. labrusca*, *V. cinerea*, *V. rupestris*, *V. acerifolia*, *V. amurensis*, *V. vulpina*, *V. aestivalis*, *V. palmata*, *V. coignetiae*, *Ampelopsis glandulosa* var. *brevipedunculata*, *V*. ×*andersonii*, *V*. ×*champinii*, *V*. ×*doaniana*, *V*. ×*novae-angliae*, and 13 vines with unassigned identity (indicated as *Vitis* spp.). To help simplify visualization, only the six most well-sampled species (*V. riparia*, *V. acerifolia*, *V. labrusca*, *V. amurensis*, *V. rupestris*, and *V. cinerea*) are separately indicated in plots and the remainder are grouped as “Other” (**Fig. 1**).

We reject the European common names for Passifloraceae species that romanticize colonialism and force Christianity over the Indigenous cultures that first described these species <u>(Dwyer et al. 2022)</u>, and instead use the Hispanicized name *maracuyá*, derived from Old Tupí or Guaraní. Maracuyá data is previously described in Otoni and Chitwood <u>(Chitwood and Otoni 2017b, 2017a)</u>. Original leaf scans <u>(Chitwood and Otoni 2016)</u> and landmark data <u>(Chitwood [2016] 2023)</u> are both publicly available. More than 3,300 leaves were collected from up to 40 species keeping track of the node, counting from the growing tip of the vine, they were collected from. Species have been clustered into Morphotypes A-G morphologically (not phylogenetically) as described in Otoni and Chitwood <u>(Chitwood and Otoni 2017b)</u> as follows: Morphotype A – *Passiflora coriacea*, *P. misera*; Morphotype B – *P. biflora*, *P. capsularis*, *P. micropetala*, *P. organensis*, *P. pohlii*, *P. rubra*, *P. tricuspis*; Morphotype C – *P. caerulea*, *P. cincinnata*, *P. edmundoi*, *P. gibertii*, *P. hatschbachii*, *P. kermesina*, *P. mollissima*, *P. setacea*, *P. suberosa*, *P. tenuifila*; Morphotype D – *P. amethystina*, *P. foetida*, *P. gracilis*, *P. morifolia*; Morphotype E – *P. actinia*, *P. miersii*, *P. sidifolia*, *P. triloba*; Morphotype F – *P. alata*, *P. edulis*, *P. ligularis*, *P. nitida*, *P. racemosa*, *P. villosa*; Morphotype G – *P. coccinea*, *P. cristalina*, *P. galbana*, *P. malacophylla*, *P. maliformis*, *P. miniata*, *P. mucronata* (**Fig. 1**).

### Mapper algorithm

Each leaf in the dataset is represented by 15 *(x,y)*-coordinate pairs, one pair denoting the location of each landmark (**Fig. 2B**). This collection of coordinates can be represented as a 30-dimensional vector, one vector for each leaf. Because direct visualization of such high-dimensional data is impossible, we use the Mapper algorithm to simplify the shape of the data into a one-dimensional graph <u>(Singh et al. 2007)</u>. For instance, Mapper was recently used to uncover patterns in gene expression in flowering plants which PCA failed to distinguish <u>(Palande et al. 2022)</u>. The Mapper algorithm works by creating groups of nearby points, then using these groups as a basis for the graph structure.

**Figure 2.**
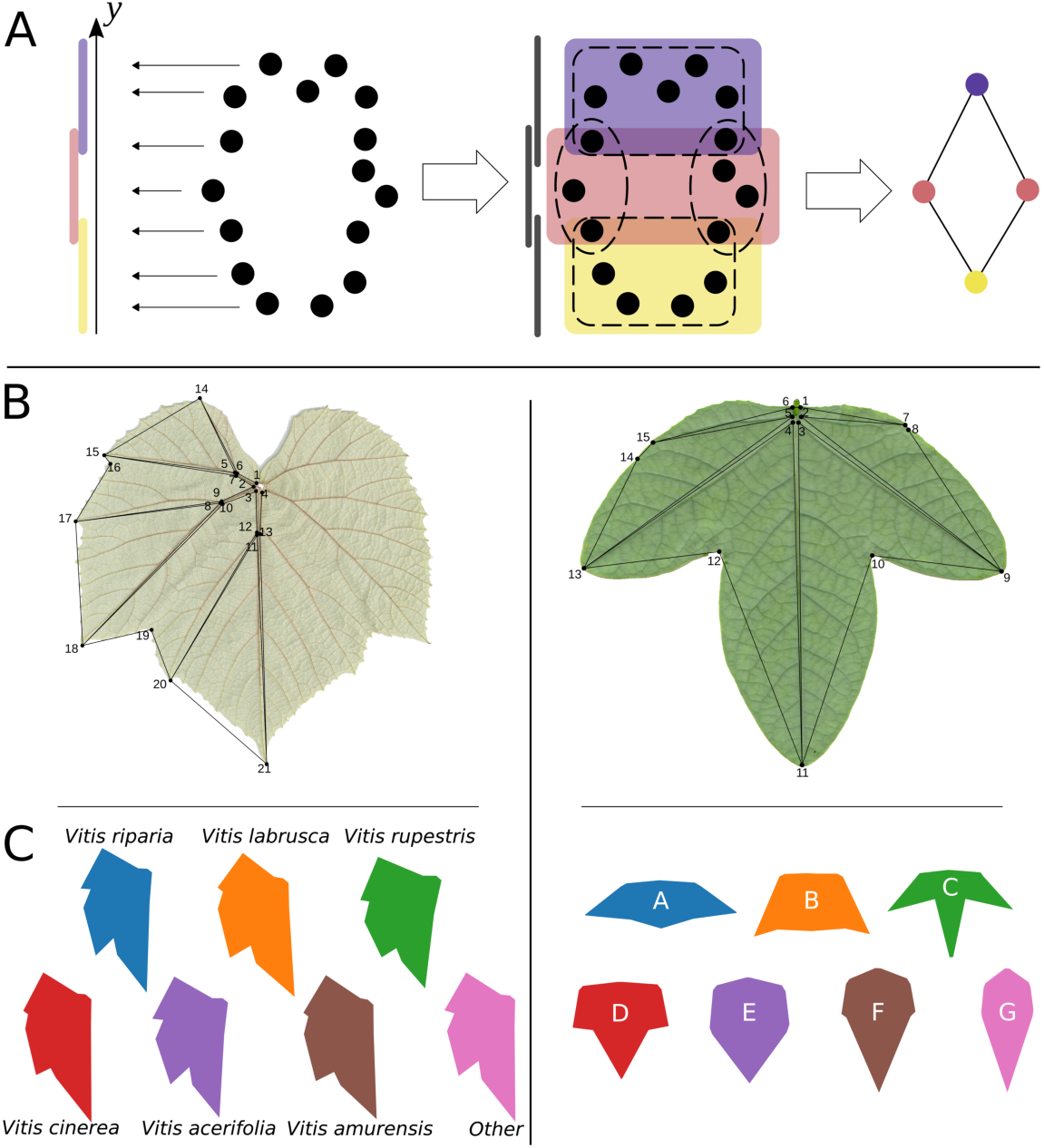
Mapper construction, leaf landmarks, and representative examples from grapevine and maracuyá morphotypes. (A) Generalized overview of Mapper graph construction and lens function selection (see **Materials and Methods** for details). Each data point has a distance from every other and is assigned a real number value through a lens function, *y* in this case (left). Points are binned by overlapping cover intervals across the lens function, clustered into nodes, and edges placed between nodes spanning cover intervals with shared data points to create a graph (right). **(B)** Landmarks used for grapevine (left) and maracuyá (right). **(C)** Averaged leaf shapes representing grapevine species (left) and maracuyá morphotypes (right).

Let X be a point cloud, and consider a *lens function* f: X → ℝ. While the Mapper algorithm is defined for more general maps, in this work, we only consider maps f: X → ℝ. The Mapper algorithm consists of the following steps:

1. Construct an open cover {U_α_} of f(X). In practice, each U_α_ is an interval in ℝ, with endpoints determined by the user-specified number of intervals and their overlap.
2. Cluster the points within each open f^-1^(U_α_).
3. The Mapper graph is the one-skeleton of the nerve of the clustering from the previous step; each cluster corresponds to a vertex in the Mapper graph, with two vertices being connected if there is a point in each of their respective clusters.

The mathematical description above can be described qualitatively, as follows. We think of our data as a point cloud, where the coordinates of each point, or leaf, are determined by the 30- dimensional vector containing the coordinates of each landmark. To use the Mapper algorithm, we must first have a notion of “distance” from any leaf to any other leaf. We choose to use the correlation distance, defined as 1-r, where r is the Pearson correlation coefficient, because in this semi-metric, leaf vectors with high correlation between their landmarks will have a distance near zero (**Fig. 2A**).

At a high level, the first step in the Mapper algorithm is to group points with similar points. There are many ways to do this, but the implementation of the Mapper algorithm we use follows a specific protocol: first, we perform a dimensionality reduction step by mapping each leaf to a value determined by a lens function. This lens function is user-defined and is determined by which aspect of the data is the topic of study. In our work, we make use of two lens functions: the first PCA component, and heteroblasty.

Now that the data has been mapped to one dimension, we cover these projected data points with a set of overlapping intervals. The open cover used in the Mapper algorithm is arbitrary and affects the structure of the resulting graph. Heuristically, coarser covers lead to graphs with fewer vertices and finer covers result in graphs with more vertices. Similarly, higher amounts of overlap between cover intervals generally result in graphs with more edges between vertices. In this work, we hand-tune the open cover to obtain a graph that is sufficiently detailed while still being human-readable. Once the intervals in the open cover have been defined, we use these intervals to group points in the original data set: two points are in the same group if their projected images are in the same interval.

The next step is to build the Mapper graph from these groupings. Inside of each group, we perform a clustering algorithm. We use DBSCAN <u>(Ester et al. 1996)</u> since it does not require the user to select the number of clusters *a priori* and is robust to outliers. We construct the Mapper graph as follows: each cluster becomes a vertex in the Mapper graph. Vertices are connected if both vertices contain a common data point; this is possible because one data point may be in multiple intervals. The resulting graph is a one-dimensional Mapper graph.

## RESULTS

### Mapper resolves relationships within grapevine and maracuyá better than PCA

Leaves are high-dimensional shapes derived from the integration of developmental, environmental, and evolutionary forces (**Fig. 1**). To make sense of high-dimensional datasets, traditional data analysis often relies on Principal Component Analysis (PCA). We therefore revisited morphological datasets of grapevine and maracuyá to see whether dimension reduction via PCA could provide novel insights into the relationships between these complex and variable leaf shapes (**Figs. 1, 3A-B**). PCA plots of maracuyá leaves partially separated the seven morphotypes but still retained significant overlap near the center of the graph (**Fig. 3B**). Similar analyses using grapevine leaves were even less successful at distinguishing individual species, with all points essentially clustering into one large group (**Fig. 3A**). Thus, while PCA plots provide some separation between groups, they are highly variable and have considerable overlap, obscuring relationships across genetic or developmental factors.

**Figure 3.**
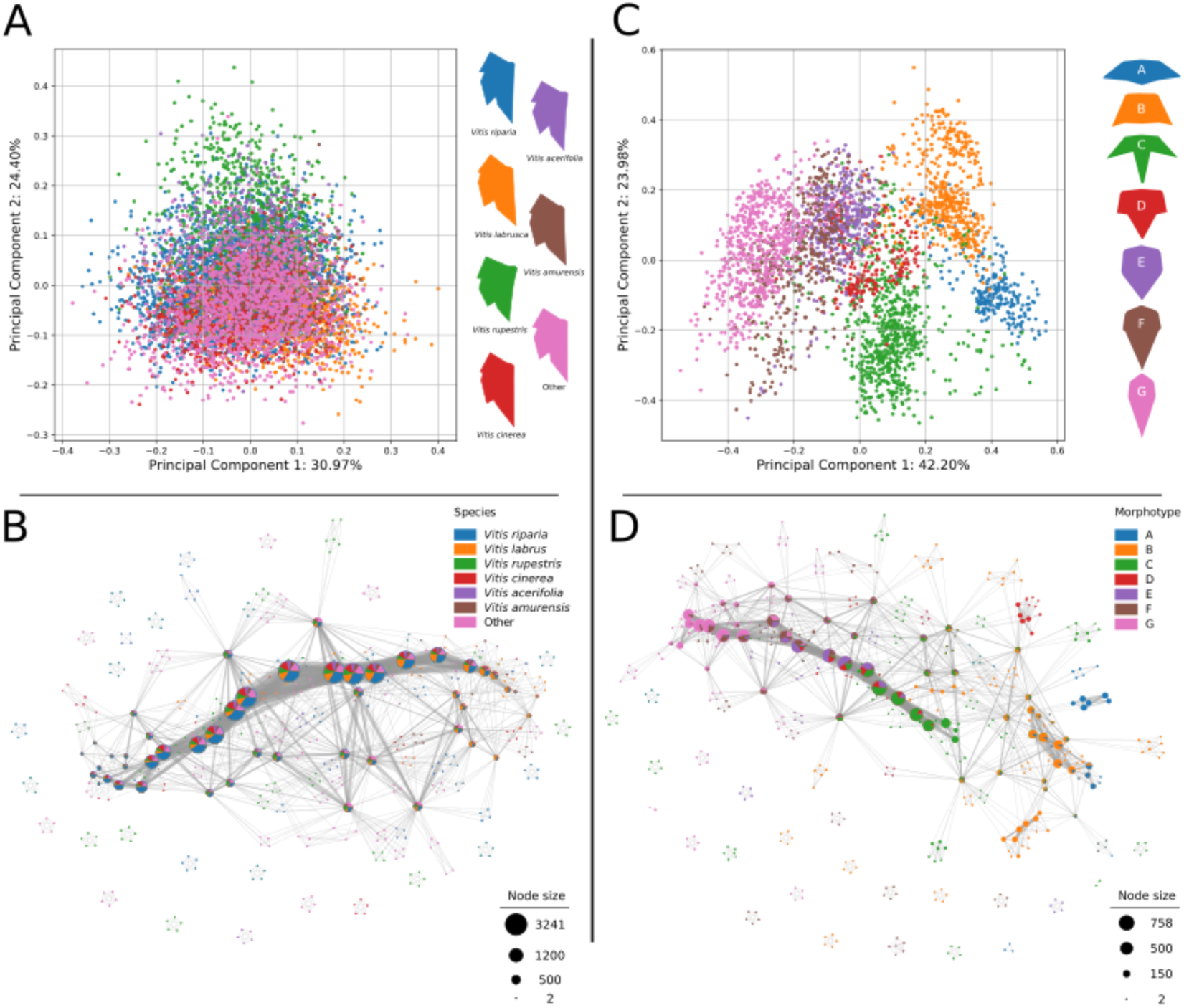
Principal component analyses (PCA) and PC1-derived Mapper graphs of grapevine and maracuyá morphotypes. (A) PCA fails to clearly distinguish grapevine morphotypes. **(B)** A Mapper graph using PC1 as a lens retains the underlying structure of the PCA plot and better resolves membership within each group. **(C)** PCA resolves maracuyá morphotypes better than grapevine but shows substantial overlap near the center of the plot. **(D)** A Mapper graph using PC1 as a lens retains the underlying structure of the PCA plot and better resolves membership within each group.

As an alternate approach, we turned to the Mapper algorithm, a method that has successfully identified novel relationships within other high-dimensional datasets <u>(Palande et al. 2022; Rathore et al. 2023; Rizvi et al. 2017)</u>. A key advantage of Mapper is its ability to interrogate datasets using several different lenses, revealing structure and relationships from specific, mathematically-defined perspectives that are hidden from traditional approaches (see **Materials and Methods**). First, we tested whether Mapper could recapitulate the relationships between grapevine and maracuyá species identified by PCA. We did this by using Mapper to visualize the structure of the data from the perspective of PC1. By representing vertices in the Mapper graph with pie charts denoting their composition, we were able to better resolve group membership within each vertex for both maracuyá and grapevine (**Figs. 3B, 3D**). Further, because PC1 was chosen as the lens, the structure of each Mapper graph retains the underlying structure of its respective PCA plot. For instance, the branches of the maracuyá Mapper correspond to the clusters of its PCA both in relative position and in membership (**Fig. 3D**). By contrast, vertices of the grapevine Mapper do not show clear differences in membership, consistent with the single clustered nature of its PCA (**Fig. 3B**). Thus, Mapper can recapitulate and improve on results from PCA, and is positioned to identify new relationships that this traditional approach cannot.

### Mapper graphs reveal conserved developmental programs in grapevine and maracuyá

The maracuyá and grapevine datasets contain several replicates of leaves collected from every node along growing vines. This presents the opportunity to explore developmental questions such as heteroblasty and ontogeny. Further, as the datasets contain several species per genus, these questions can also be examined in an evolutionary context. To begin to address the interplay between evolution, development, and leaf shape, we assigned a relative heteroblasty value to each leaf based on its position, ranging from zero (tip of vine) to one (base of vine). Recoloring the PCA plot for grapevine using these values revealed juvenile leaves cluster near the bottom left of the plot while adult leaves cluster towards the top right (**Fig. 4A, left**). This suggests a relationship between heteroblasty and both PC1 and PC2, however this relationship is obscured by the many overlapping points on the plot. Recoloring the PCA plot for maracuyá revealed a more complicated distribution, with juvenile and adult leaves displaying a significant amount of overlap and no obvious pattern (**Fig. 4, right**). Heteroblastic trends within and between species are thus not clearly delineated using traditional analyses like PCA.

**Figure 4.**
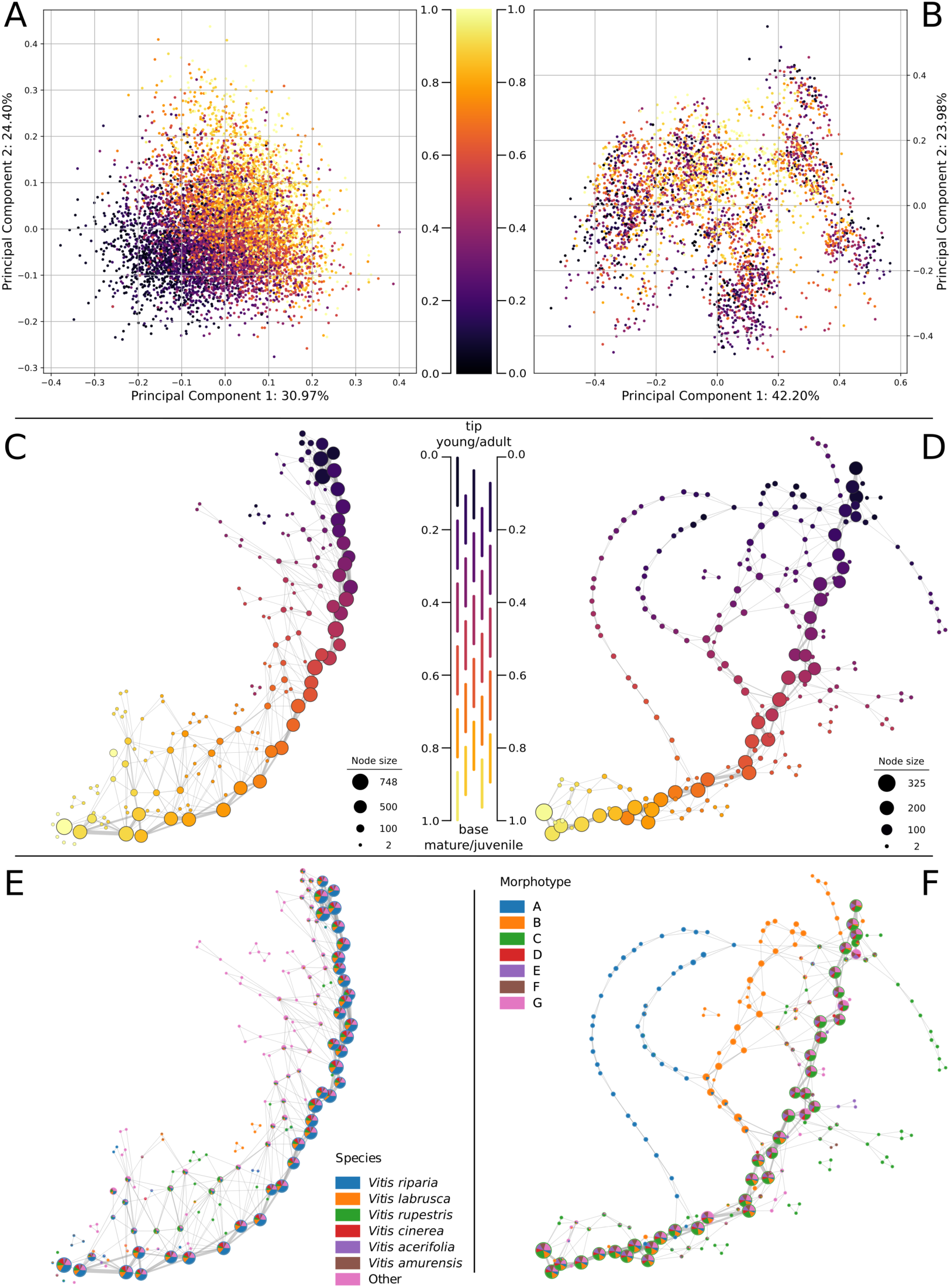
Grapevine and maracuyá species share a core conserved heteroblasty program. (A) Recoloring of the grapevine PCA by heteroblasty values suggests a relationship between heteroblasty and both PC1 and PC2. **(B)**. Recoloring the PCA plot for maracuyá yielded no obvious relationships. **(C,E)** A Mapper graph using heteroblasty as a lens reveals a strong central spine in grapevine shared by all morphotypes. **(D,F)** A Mapper graph using heteroblasty as a lens reveals a strong central spine in maracuyá shared by most morphotypes. Two notable exceptions are morphotypes A and B which diverge near the middle of the leaf series before rejoining the central spine.

To better identify these hidden relationships, we constructed Mapper graphs for maracuyá and grapevine using heteroblasty as a lens (**Fig. 4C,D**). Vertices within the resulting Mapper graphs were colored by the average heteroblasty value of each leaf in the vertex. One prediction of the recolored PCA plots is that grapevine should have limited branching, given its relatively clear continuum of heteroblasty values along PC1 and PC2 (**Fig. 4A**). Supporting this, the Mapper graph for grapevine consists of a strong central spine, with clear transitions between juvenile and adult leaves, and limited branching (**Fig. 4C, 4E**). A second prediction is that maracuyá should have a much more complex structure. Indeed, given their strikingly different leaf morphologies (**Fig. 1**), each morphotype could conceivably have its own heteroblastic trajectory (**Fig. 4C**). Surprisingly, the Mapper graph for maracuyá also has a strong central spine (**Fig. 4D**). As the Mapper algorithm detects shape changes between concurrent nodes, this suggests that most maracuyá species share a core, deeply conserved heteroblasty program despite the markedly different appearances of their leaves (**Fig. 1)**. Two interesting exceptions to this are morphotypes A and B which branch off from the central spine at two distinct points (**Figs. 4D, 4F**). These morphotypes are members of the phylogenetically distant *Decaloba* group (Muschner et al., 2003; Krosnick et al., 2013), suggesting the evolution of a distinct heteroblasty program in this group. Interestingly, both morphotypes eventually rejoin the main spine, suggesting the beginning and ending of this distinct program is shared by the entire maracuyá family.

The power of Mapper is its ability to visualize hidden structure within high-dimensional datasets which is then presented as an abstract graph composed of edges and vertices. However, to fully understand how shape changes across a given lens (in this case heteroblasty), it is helpful to relate this graphical representation back to the real shapes that drove its construction. To do this, we first extracted the primary structure of the Mapper graphs by highlighting their central spines and branching structures (**Fig. 5**). Vertices were then replaced by leaf outlines which represent the average shape of all leaves within a given vertex. The resulting morphospaces provide a tangible illustration of how leaf shape changes along grapevine and maracuyá shoots. For instance, leaf outlines along the central spine of the grapevine Mapper show little deviation, as expected (**Fig. 5A**). By contrast, the representative outlines along the central spine of maracuyá are highly variable indicating no single morphotype dominates the morphospace (**Fig. 5B**). Thus, despite members of the maracuyá group displaying striking differences in leaf shapes (**Fig. 1**), the way their leaves change from node to node is deeply conserved. Exceptions to this – morphotypes A and B – are also visible as branches emerging from different points along the main spine. Whether their evolutionarily distinct nature plays into this branching is an open question.

**Figure 5.**
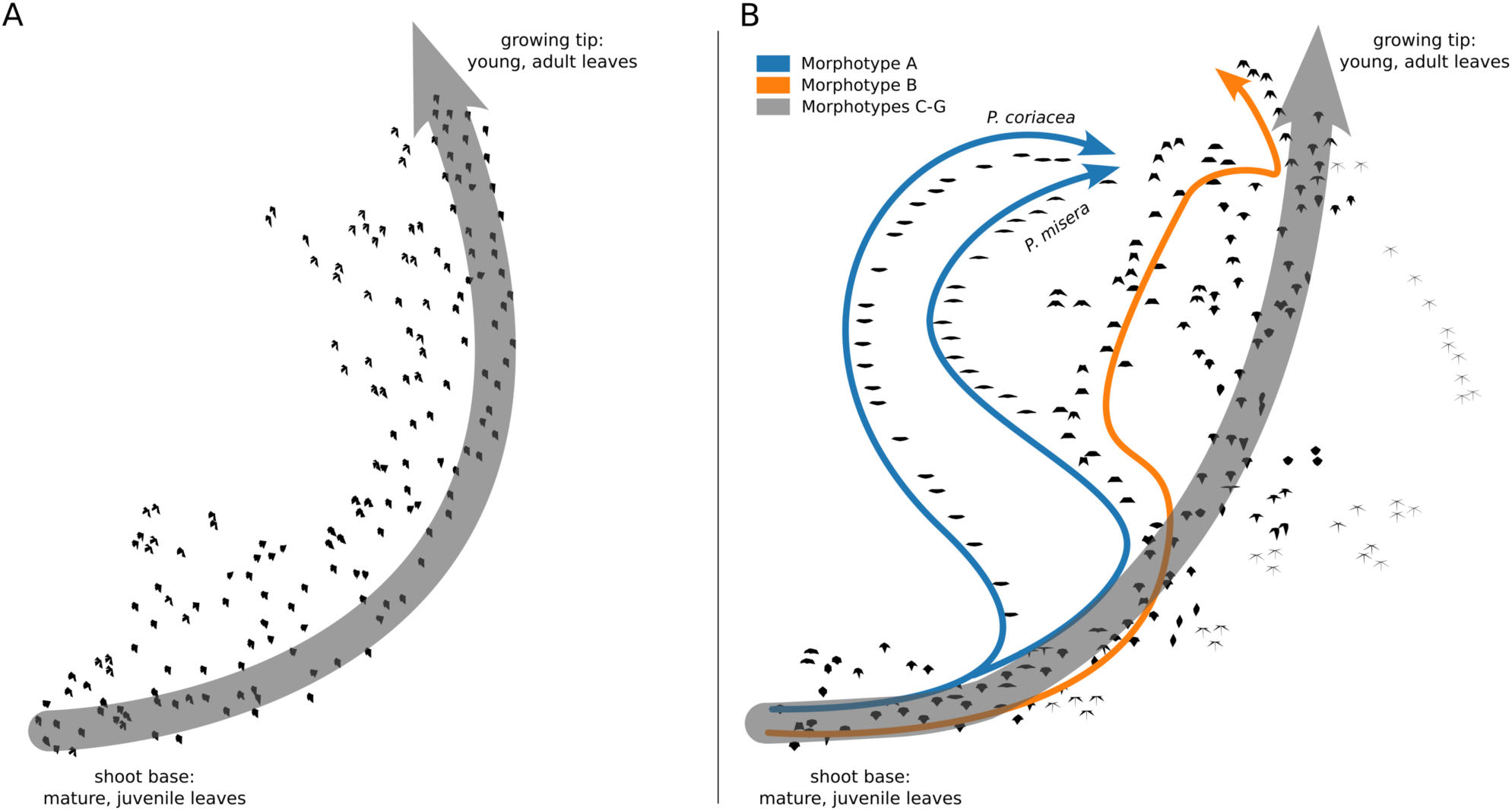
Subway map of grapevine and maracuyá Mapper graphs. (A) Primary structure of the grapevine Mapper graph with vertices represented by the average leaf shape within a given vertex. **(B)** Primary structure of the maracuyá Mapper graph with vertices represented by the average leaf shape within a given vertex. No single morphotype dominates each vertex demonstrating that Mapper is clustering species based on an underlying data structure and not by shape similarities. Morphotypes A and B, which display unique heteroblastic trajectories, are labeled in blue and orange, respectively.

## DISCUSSION

### A similar heteroblastic trajectory underlies most species of grapevine and maracuyá

Leaves are the product of development, environmental, and evolutionary forces, and their shapes vary dramatically both within plants of a given species and between species of a given family. One example of these forces is a process termed heteroblasty, in which leaves of different shapes are generated from a single plant in an age-dependent manner (Kerstetter & Poethig, 1998). In our study, the leaves of the maracuyá family provide a particularly striking illustration of this, though subtle changes can be seen in grapevine, as well (**Fig. 1**). Modulation of a core heteroblasty program is thus a logical potential explanation for the species-to-species divergence of leaf morphologies seen between these two families (Cartolano et al., 2015; Willman and Poething, 2011). Surprisingly, a Mapper algorithm using a heteroblastic lens revealed that most species, including those in the maracuyá family, have a nearly identical trajectory despite their highly divergent leaf shapes (**Fig. 4**). Topological data analysis thus allows the contribution of a given process – in this case heteroblasty – to be isolated from the other forces guiding leaf morphogenesis. One potential caveat concerns leaf measurements from the tip of the growing vines. As these leaves are not fully mature, node-to-node shape changes are likely influenced by both ontogenetic and heteroblastic pathways. Importantly, biologically relevant relationships in this region were still identified and preserved by the Mapper algorithm, supporting the robustness of this approach.

### Modulating leaf shape by coupling slowly and quickly evolving processes

If not heteroblasty, then what other forces or factors might underlie the species-to-species differences in leaf shape seen in these datasets? Multiple molecular pathways intersect to control leaf shape, and there is no shortage of candidates (reviewed in Chitwood and Sinha, 2016). For instance, leaf complexity is regulated by several transcription factors that in some cases operate independently of heteroblasty. The homeobox gene LATE MERISTEM IDENTITY1 (LMI1) / REDUCED COMPLEXITY (RCO), for example, drives evolutionary differences in leaf shape between species within Brassicaceae (Saddic et al., 2006 Development; Vlad et al., 2014 Science) and between varieties of cotton (Andres et al., 2017). Similarly, mutations in KNOX genes can confer evolutionarily labile effects on leaf complexity between closely related species of tomato (Kimura et al., 2008 Curr Biol). Regardless of the specific factor, a plausible explanation for species-specific differences in maracuyá might be evolutionarily labile effects of genes regulating leaf morphology, similar to those described above, layered on more slowly evolving developmental pathways such as auxin signaling <u>(Koenig et al. 2009)</u>, adaxial-abaxial patterning <u>(Husbands et al. 2009)</u>, proximal-distal patterning <u>(Das Gupta and Nath 2015)</u>, and miR156-miR172-mediated heteroblasty (Wu et al., 2009 Cell), the effects of which are conserved in maracuyá (<u>Silva et al. 2019)</u>. Gene regulatory networks distinct from conserved pathways regulating heteroblastic transitions across the flowering plants would be expected to confer different effects on leaf shape and might explain the independence of these two processes we observe at a morphological level.

### A ‘reverse hourglass’ of heteroblasty in select maracuyá morphotypes

Two interesting exceptions emerged from our analyses of maracuyá leaf shape. Whereas all maracuyá species cluster at the growing tip and base of their shoots, morphotypes A and B in the *Decaloba* group diverge in the middle of the leaf series (**Figs. 4, 5**). One useful metaphor to describe this relates to the hourglass model of gene expression which posits that gene expression programs are more conserved near the middle of embryogenesis than the beginning and end <u>(Quint et al. 2012)</u>. In the case of the *Decaloba* group, our data reveal a “reverse hourglass” effect, with similarity highest near the beginning and end of the leaf series, and lowest near the middle. This suggests that these species deploy the standard maracuyá heteroblasty program at their tip and base but evolved a unique program near the middle of their leaf series. Alternatively, the core heteroblasty program may be conserved throughout, but heteroblasty-independent effects exert a particularly strong influence in this region in these morphotypes. Genetic and molecular assays could be used to distinguish between these scenarios. Nevertheless, these findings illustrate the ability of topological data analysis to highlight biologically relevant relationships.

### Perspectives

Questions in biology are increasingly driven by large datasets comprised of ecological, morphological, and molecular measurements. These datasets contain enormous amounts of information, far more than many biologists appreciate. For instance, even a single leaf on a growing plant is defined by multiple identities and contains volumes of information. Morphologically, it has length, width, and depth, and is growing along these axes at different, allometric rates. Evolutionarily, it is a member of a particular species with a defined evolutionary history. Ecologically and physiologically, it is located at specific latitude and longitude coordinates and is experiencing a local microclimate. It has also been exposed to a unique combination of environmental conditions over its lifetime. At the granular level, it is comprised of hundreds of cells clustering into distinct cell types such as the epidermis, vasculature, and mesophyll. Each of these cells has its own unique molecular identity defined by chromosomes with specific combinations of epigenetic marks, a transcriptome, a proteome, and a metabolome. Note that these are merely a subset of ways that one could define a single leaf. The central challenge of the field is to make sense of this enormous amount of information. Topological data analysis offers a simple, flexible, and powerful way to meet this challenge, allowing different underlying data structures arising from mathematically-defined perspectives of the same data to be visualized. Moving forward, phenotypic and molecular data structures from the same samples could be directly compared to each other or modeled after each other, discerning previously confounded mechanisms and ultimately permitting the development of predictive models of complex phenotypes from underlying molecular and genetic signatures. Topological data analysis is thus poised to provide transformative insights into a wide range of biological questions.

## DATA AVAILABILITY STATEMENT

Data and code necessary to reproduce the analyses in this work can be found at the following link: https://github.com/sperciva/maracuya-grapevine-TDA

## ACKNOWLEDGMENTS

This work is supported by NSF awards IOS-2039489, 2310355, 2310356, and 2310357. This project was also supported by the USDA National Institute of Food and Agriculture, by Michigan State University AgBioResearch, and by Michigan State University Biochemistry & Molecular Biology Plant Biology TEAM-UP.

